# Genetic Architectures of Medical Images Revealed by Registration and Fusion of Multiple Modalities

**DOI:** 10.1101/2023.07.27.550885

**Authors:** Sam Freesun Friedman, Gemma Elyse Moran, Marianne Rakic, Anthony Phillipakis

**Affiliations:** Broad Institute of MIT and Harvard; MIT

## Abstract

The advent of biobanks with vast quantities of medical imaging and paired genetic measurements creates huge opportunities for a new generation of genotype-phenotype association studies. However, disentangling biological signals from the many sources of bias and artifacts remains difficult. Using diverse types of medical imaging (i.e. MRIs, ECGs and DXAs), we develop registered and cross-modal generative models. In all cases, we show how registration, both spatial and temporal, guided by domain knowledge or learned de novo, uncovers rich biological information. Remarkably, our findings demonstrate that even extremely lossy transformations, such as registering images onto a single 1D curve (e.g. a circle), can yield robust signals. Conversely, we demonstrate that increasing data dimensionality by integrating multiple modalities can also result in richer representations. Through genome- and phenome-wide association studies (GWAS and PheWAS) of learned embeddings, we uncover significantly more associations with registered and fused modalities than with equivalently trained and sized representations learned from native coordinate spaces. Our findings systematically reveal the crucial role registration plays in enhancing the characterization of physiological states across a broad range of medical imaging data types.

## Introduction

The concept of registration – mathematical maps between coordinate systems – has ancient roots, dating back at least as far as the ancient Egyptians(Imhausen 2020). They surveyed land with regularly knotted ropes which were gathered, physically implementing isomorphic scaling(Barnard 2008). The Cartesian grid put the idea on solid mathematical footing and the problem has attracted interest and innovation ever since(Descartes 1886; Goshtasby 1988; Friedman and Stamos 2013; Ardeshir Goshtasby 2005).

Given a medical image, one way to understand the similarity between different individuals is to align them to a common prototypical example of the data, called an atlas. The common reference can be a prototypical example of the data (i.e. an atlas or template), or an idealized parameterization of an anatomical feature. Here, we show that medical image registration has a profound impact on the representations that deep neural networks learn, vastly increasing the encoded biological signal. This is demonstrated across a broad range of registration methods from hand-crafted templates to learned non-linear deformation fields, and a correspondingly broad range of medical image from electro-temporal 1D ECGs to spatial 2D DXAs, 3D brain MRIs and spatio-temporal 3D cardiac MRIs.

At a high level, registration helps models target the biological variance of interest. Variation from non-biological sources is common in medical images: for example, MRIs may encode technical or environmental sources of variation, such as operator dependence or the position of a patient in the scanner. Such nuisance variation is often referred to as ‘spurious correlation’. In the supervised learning setting, there is a large literature on how to adjust for such spurious correlations. These approaches include: leveraging data augmentations(Puli et al. 2022); adjustment based on an assumed causal model(Wang and Jordan 2021); adjustment using auxiliary labels(Makar et al. 28--30 Mar 2022); and regularizing prediction functions across different domains(Nguyen et al. 2021). The unsupervised learning setting, meanwhile, presents an even greater challenge as there is no outcome variable to help focus the model on relevant signals. To ameliorate this issue, we demonstrate how domain-specific registrations can be used to learn biologically-informed representations.

The recent availability of large-scale multimodal measurements in biobanks(Sudlow et al. 2015; Bycroft et al. 2018) provides the opportunity to systematically study how registration impacts representations of physiology. Because biobanks often assess individuals with many different modalities they also allow us to investigate how fusion of different image types can lead to new insights. Consistent with our observations, a line of recent works utilize contrastive encoders to learn joint representations of multimodal data, including natural images and captions in computer vision(Ramesh et al. 2022; Radford et al. 2021), and paired clinical measurements(Diamant et al. 2022; Vu et al. 06--07 Aug 2021). Contrastive fusion can be thought of as a learned registration between distinct modalities and, like simpler registration techniques, it too reveals more genetic and phenotypic associations(Radhakrishnan et al. 2023).

### Related Work

#### Anatomical Registrations

Many anatomical features have been used for registration. For example, 3D anatomical atlases of the brain are used to register brain MRIs(Tzourio-Mazoyer et al. 2002), while the cardiac cycle of a single heartbeat can serve as the anatomical landmark for ECGs and cardiac MRIs(Carreiras et al., n.d.). Similarly, the temporal mapping of ECGs to a single cardiac cycle is achieved through template matching of the QRS complex, followed by alignment, scaling and median computation of each waveform from the full 10 seconds of the resting ECG tracing(Carreiras et al., n.d.). DXA scans can be registered with rigid homomorphic mappings to an exemplar individual(Ardeshir Goshtasby 2005). Figure 1 provides visual examples of anatomical registration by template matching and atlas alignment.

**Figure 1:**
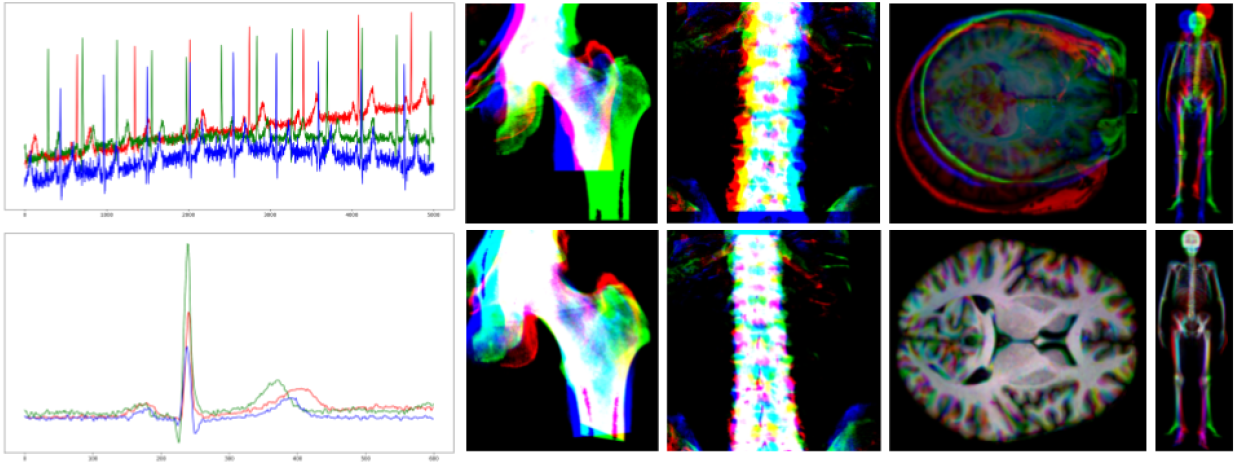
Examples of 3 individuals overlaid as color channels before and after registration. The top row shows the original modality and the bottom row shows the registered version. From left to right the modalites shown are the resting ECG, DXA 5 (hip), DXA 2 (lumbar spine), T1 brain MRI, and DXA 11 (whole-body skeletal).

#### Learned Registrations

Classical methods solve a new optimization problem for every registration pair, and they are therefore computationally expensive(Vercauteren et al. 2009; Avants et al. 2011; Shen 2007; Thirion 1998). Some deep learning methods propose to learn the alignment between the medical image and reference instead(Balakrishnan et al. 2019; Yu et al. 2022; Yang et al. 2017; de Vos et al. 2017); supervised methods propose to train the network and compare the output to pre-computed alignments, while unsupervised methods train their networks by learning a mapping between the image and reference (deformation field) and comparing how well the aligned image matches the reference(Jaderberg et al. 2015). VoxelMorph(Balakrishnan et al. 2019), for example, uses convolutions and spatial transformations together with a UNet architecture to learn a deformation field for each new image/reference pair. It learns amortized registration by jointly maximizing agreement between the aligned image and the reference, as well as transformation smoothness(Balakrishnan et al. 2019). While deformation fields can preserve or even increase parameterization of a modality, many registration techniques reduce dimensionality. Taking reduction to its logical extreme, DeepCycle uses a single-parameter autoencoder and the inductive bias of periodicity, to register single cell RNA expression data to the mitotic cell-cycle(Riba et al. 2022). Recent multimodal fusion methods, like DropFuse, use contrastive cross-modal learning to register different modalities into a single 256-dimension latent space, also greatly reducing data size(Radhakrishnan et al. 2023).

#### Genetic Analysis of Medical Images

The advent of large genotyped Biobanks enables whole genome association testing with traits derived from medical images such as MRIs, retinal images and ECGs(Elliott et al. 2018; Zekavat et al. 2022; Haas et al. 2021). Active research has extended these genomic analyses from traits to spaces, performing the association tests in unsupervised ways. For example, GWAS of each dimension in a variational autoencoder, or the principal components of that encoding or even with ECG voltages from medians over every millisecond of input(Verweij et al. 2020; Yun et al. 2023; Xie et al. 2022; Kirchler et al. 2022).

Building on this work, we demonstrate how across this broad and diverse sample of methods and modalities, registration uniformly results in richer phenotypic and genotypic associations.

## Methods

We train modality-specific DenseNet-style(Iandola et al. 2014) convolutional encoders and decoders to reconstruct both registered and unregistered medical images from latent space bottlenecks. Table 1 details the registration techniques considered, the modalities they are applied to and whether the registration implementation is learned from scratch, optimized from a known transformation (homeomorphic or warp field), or hard-coded. The learned registrations are between individuals via deformation fields (VoxelMorph), with the inductive bias of periodicity (DeepCycle) or contrastively across modality (DropFuse), see Figure 2. These previously described models are briefly summarized below.

**Table 1:** Modalities and Registrations. Each modality, its original shape, shape after registration, the resulting reduction in dimensionality, the method used to register the modality, and the way the parameters of the registering transformation are derived (i.e. hard-coded with domain knowledge, optimized parameters of a known transformation or learned a new transformation via neural net approximation).

| Modality | N | Original Shape | Registered Shape | Reduction | Registration Method | Parameter Tuning |
| --- | --- | --- | --- | --- | --- | --- |
| ECG | 42k | 5000, 12 | 600, 12 | 8x | QRS Peak Align(Carreiras et al., n.d.) | Hard-coded |
| DXA | 39k | 928, 352 | 928, 352 | 1x | Homeomorphic(Bradski 2000) | Optimized |
| DXA | 39k | 928, 352 | 928, 352 | 1x | VoxelMorph(Balakrishnan et al. 2019) | Learned |
| Brain MRI | 44k | 216, 256, 216 | 182, 218, 182 | 1.2x | FreeSurfer(Fischl 2012) | Optimized |
| Cardiac MRI | 45k | 96, 96, 50 | 96, 96, 50 | 1x | VoxelMorph(Balakrishnan et al. 2019) | Learned |
| Cardiac MRI | 45k | 96, 96, 50 | 50 | 9000x | DeepCycle(Riba et al. 2022) | Learned |
| Cardiac MRI + ECG | 38k | 96, 96, 50<br>600, 12 | 256 | 1800x<br>28x | DropFuse(Radhakrishnan et al. 2023) | Learned |
| DXA 11 + DXA 12 | 39k | 928, 352<br>928, 352 | 256 | 135x | DropFuse(Radhakrishnan et al. 2023) | Learned |
| DXA 2 + DXA 5 | 39k | 768, 768<br>768, 768 | 256 | 2300x | DropFuse(Radhakrishnan et al. 2023) | Learned |

**Figure 2:**
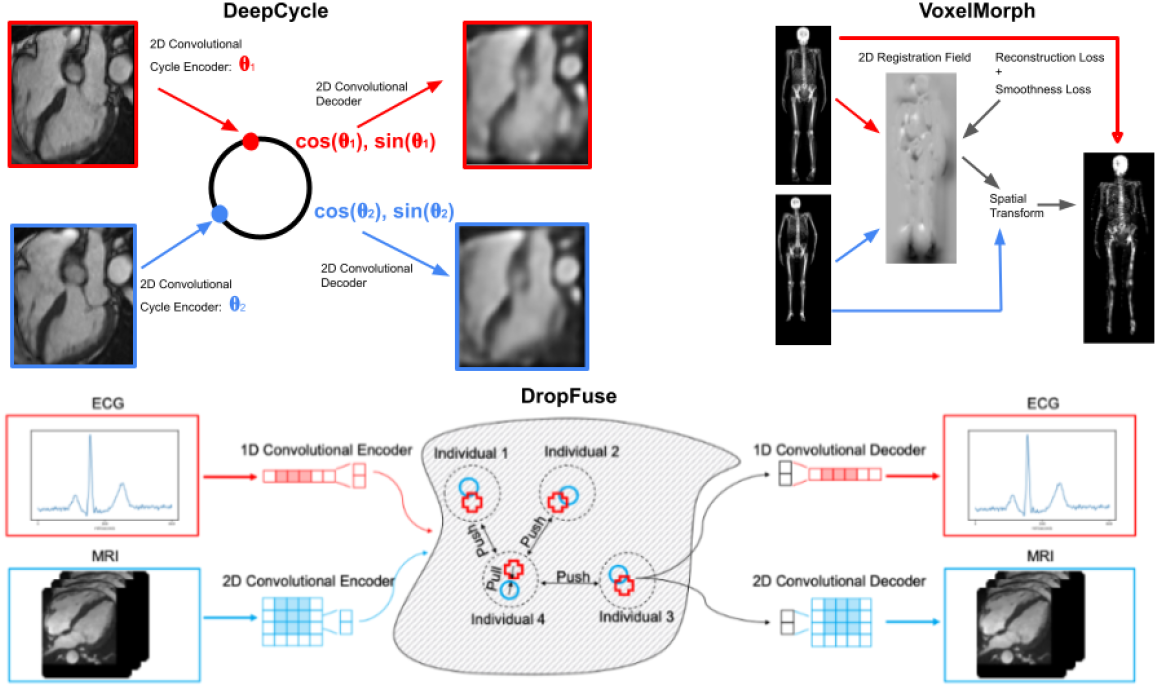
Learned Registration Methods. Three deep-learning methods used to register medical images. DeepCyle registers MRI frames to the cardiac cycle by encoding each from with a single parameter, θ, VoxelMorph learns spatial warps via pairwise amortized registration while retaining overall image dimensions and DropFuse uses dropout and cross-modal fusion to register multiple modalities together into a 256-dimensional latent space.

### DeepCycle

DeepCycle learns to encode data using a single-parameter latent space registered to the unit circle, demonstrating the extreme reductions possible with registration. A convolutional encoder learns a single parameter bottleneck, **θ**, which is registered onto the unit circle by computing (cos(**θ**), sin(**θ**)). A convolutional decoder then reconstructs the full size input image. This model is only trained to minimize mean-squared reconstruction error. This vastly under-parameterized representation still can generate high-fidelity reconstructions as well as generalizable and biologically informative representations, see Supplementary Video 1.

### VoxelMorph

VoxelMorph is trained to minimize both smoothness and similarity losses which respectively encode anatomical plausibility and fidelity of the learned registration. We trained VoxelMorph with 4-chamber long axis cardiac MRI cine series and smoothness loss weight of 0.5, see Supplementary Video 2. For purposes of comparison, the individual with median BMI was selected as an exemplar and all cMRI movies were VoxelMorphed to them.

### DropFuse

DropFuse is cross-modal autoencoder trained to minimize a reconstruction loss combined with a contrastive loss which ensures that paired ECG and MRI samples are mapped to nearby points in the latent space, while discordant pairs are pushed away. The encodings from each modality are fused with random dropout at each latent space coordinate. The model is trained with ECG and MRI or DXA series pairs, but inference requires only one modality to be available.

## Experiments

This work was motivated by the observation that autoencoder latent spaces trained from medical images in their native coordinate systems used much of their expressive power encoding aspects of the images of limited biological significance, like phase and baseline drifts in ECGs or limb orientation in DXAs. To quantitatively summarize the biological signal captured by a representation we aggregated a broad range of continuous and categorical phenotypes of clear biological import, including age, sex, BMI, and principal components of genetic ancestry, See Supplementary Figure 1 for the complete list of phenotypes. We then cross-validated the performance of linear and logistic regression models trained on representations from medical images before and after registration as summarized in Table 2. Improvements in area under the ROC curve and the coefficient of determination *R*^2^ are shown for all of the modalities and all of the registration techniques.

**Table 2:**
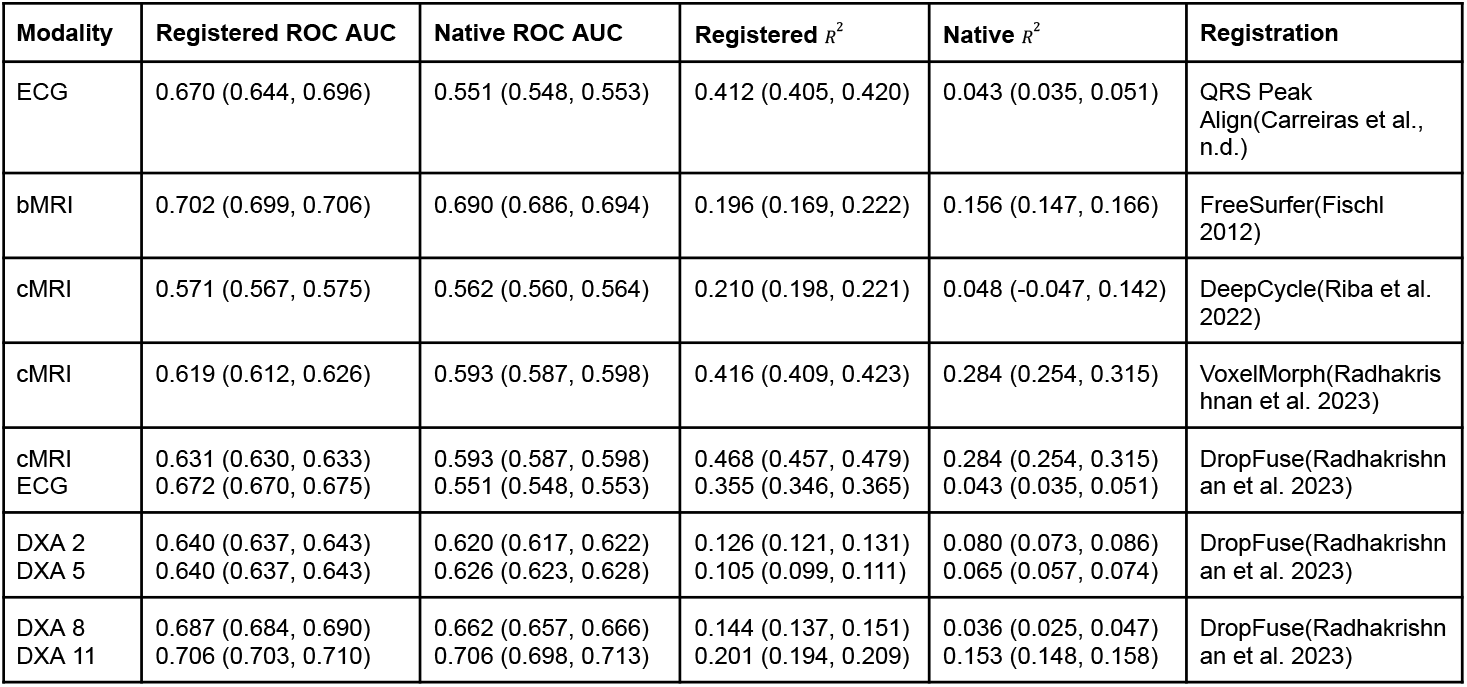
Registered modalities capture more phenotypic and genetic associations. ROC AUC and coefficient of determination are averaged across 5-fold splits of a broad range of modality-relevant tasks spanning phenotypes, diagnostics, and components of genetic ancestry. See Supplementary Figures 1-3 for performance on each of the tasks aggregated here.

Notably, even lossy registrations result in more biological signals, for example from the 5000 timepoints of the 10 second resting ECG to the 600 timepoints in the registered median waveform. Though much lower in overall dimensionality, registration greatly reduces intra-individual variations like phase and baseline drift. This frees up the latent space to “spend” its expressive capacity representing more population variation which powers downstream analyses.

Code for these experiments is available in the ML4H repository: https://github.com/broadinstitute/ml4h.

### DeepCycle Groks the Cardiac Cycle

The most lossy registration method considered (arguably the most lossy registration possible) is with the DeepCycle convolutional encoder which uses only a single parameter to encode the entire frame from cardiac MRI movie. In comparison to a convolutional autoencoder without the inductive bias of periodicity, the DeepCycle MRI representations capture more biological signal, as shown in row 3 of Table 2. Visual inspection of the DeepCycle decoder’s reconstructions reveal that the parameter, **θ**, encodes the cardiac cycle. Without the inductive bias of periodicity, encoding instead captures the exposure value of the MRI, see Supplementary Video 1. Compellingly, the DeepCyle cardiac MRI convolutional autoencoder trained using only 50 frames spanning the heart beat from a single individual, still generalizes to the larger population. Loss curves from these models consistently exhibit the recently described Grokking phenomena(Power et al. 2022). Initially, training loss decreases while validation loss increases, but after several epochs validation loss also decreases, as the model learns to better “Grok” the true data distribution, as illustrated in Figure 3.

**Figure 3:**
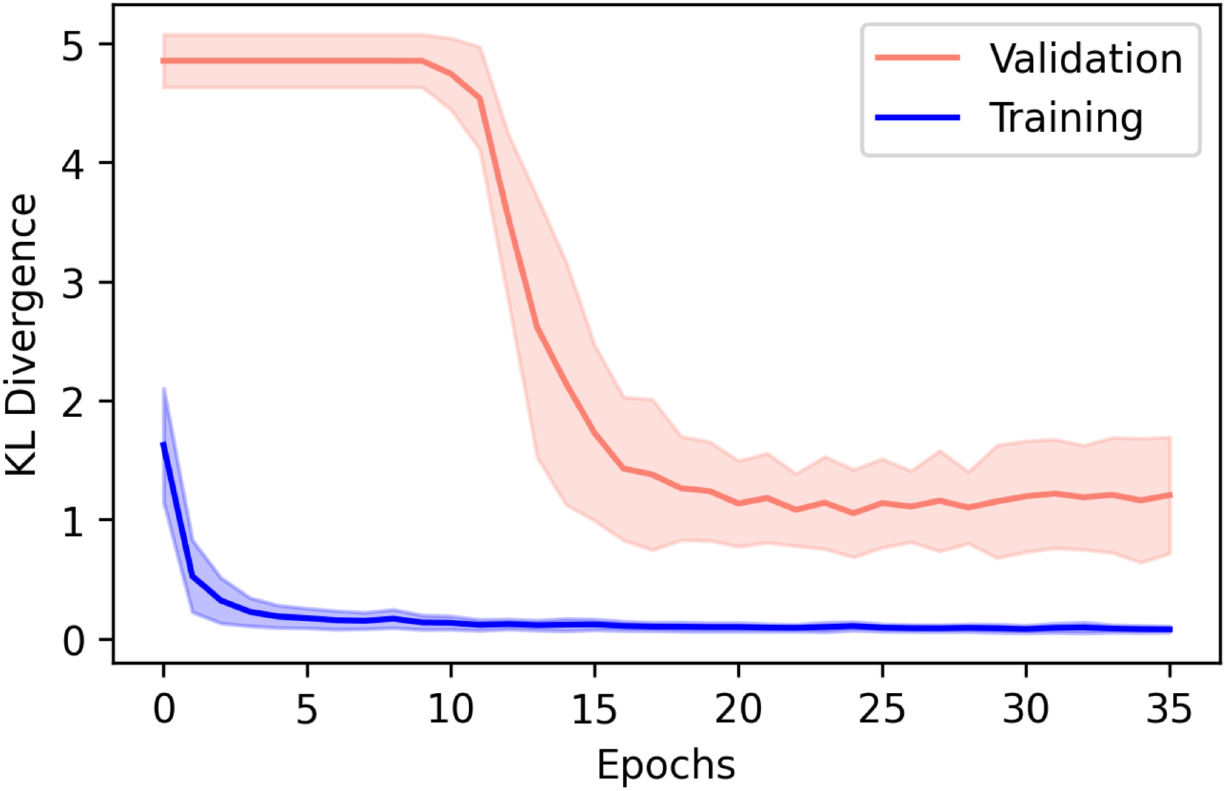
Mean (solid line) +/-one standard deviation (shaded area) of KL Divergence loss curves from 5 DeepCycle training runs with 5 different individuals and disjoint validation sets. After 10-15 epochs of training, validation loss consistently and dramatically decreases, an example of “Grokking”.

### Registered Modalities Reveal More Diagnostic Variation

Phecodes are a taxonomy of billing codes aggregated into diagnostic labels which are more reflective of true disease phenotypes(Wei et al. 2017). After determining phecode status from billing codes for the UK Biobank population, we conducted phenome wide association studies (PheWAS) before and after registration. As described in(Venn et al. 2022), with 20% of the subjects we derived latent space vectors between the centroids of individuals with and without each diagnosis. In the remaining 80% of the individuals the component in the direction of the phenotype vector was computed. Then using a logit model corrected for age, sex and race after Bonferroni corrections for multiple testing, we tabulated significant associations between phenotype vector components and each disease. The QQ plot of associations with the registered ECG latent space is shown in Figure 4. As shown in Supplementary Figures 4-6, registered latent spaces consistently identified more significant associations.

**Figure 4:**
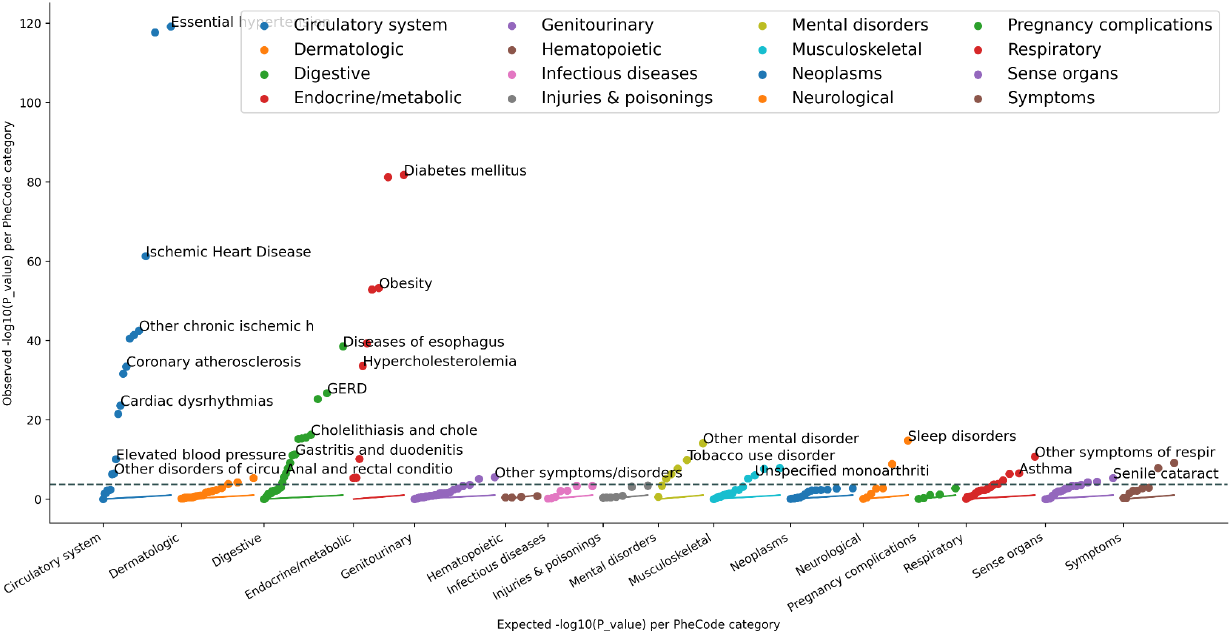
QQ plot showing − log10(P-value) for the registered ECG latent space associations with phecodes. The dotted line shows P-value threshold after Bonferroni correction for multiple testing across diagnoses

### Registered Modalities Reveal More Genetic Variation

Autoencoders trained after registration recover more genetic signals. Manhattan plots of latent space GWAS show many more loci identified by latent spaces built on the registered modalities, see Figure 5.

**Figure 5:**
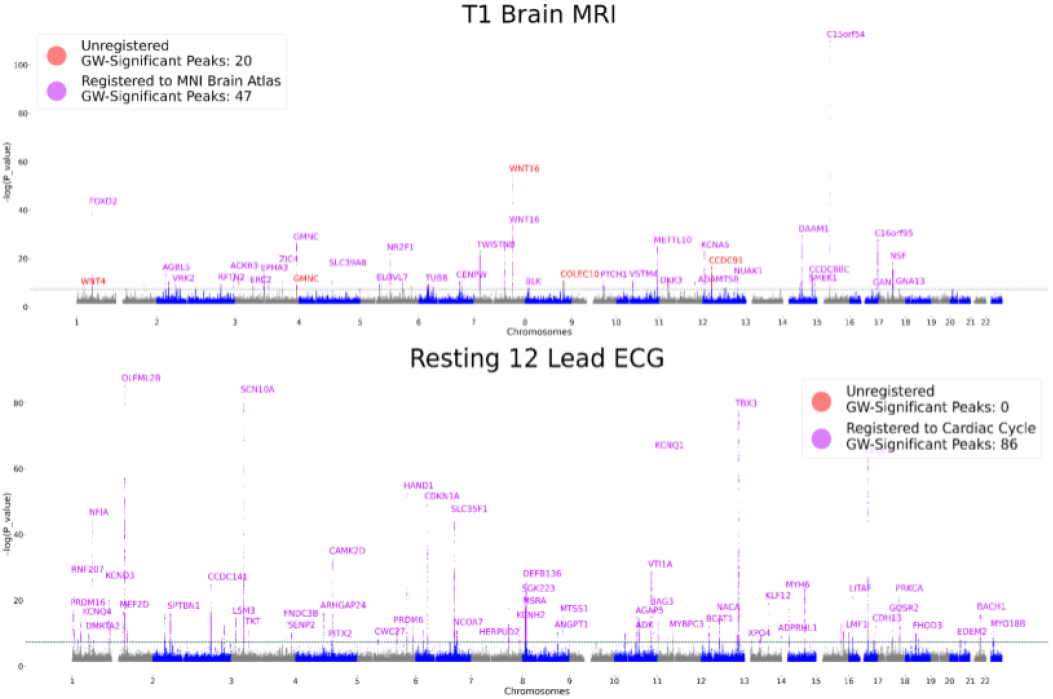
Manhattan Plots. Registration reveals more genetic associations.

We perform an “unsupervised” GWAS on autoencoder representations. For each SNP, we check if the latent space distributions of the 3 genotypes (homozygous variant, heterozygous, and homozygous reference) are separable, as quantified by Multivariate Analysis of Variance (MANOVA). Prior to performing MANOVA across the sets, we account for confounders from population stratification or batch effects, which are encoded in the embeddings by removing these sensitive features from the latent space with iterated nullspace projection(Ravfogel et al. 2020). GWAS summary statistics are deposited in the GWAS Catalog.

### Clustering SNPs in latent space elucidates genetic architectures

With this approach to genetic discovery we can cluster SNPs in the latent space to group those with similar representation and potentially similar phenotypic effects. In particular, we perform hierarchical clustering based on the direction from the mean embedding of the homozygous reference group to the mean embedding of the heterozygous and homozygous alternate groups for each lead SNP. Figure 6 contrasts latent space GWAS on anatomically registered brain MRIs demonstrating the consistency of the SNP clustering between training runs and sub-regions of the cerebellum.

**Figure 6:**
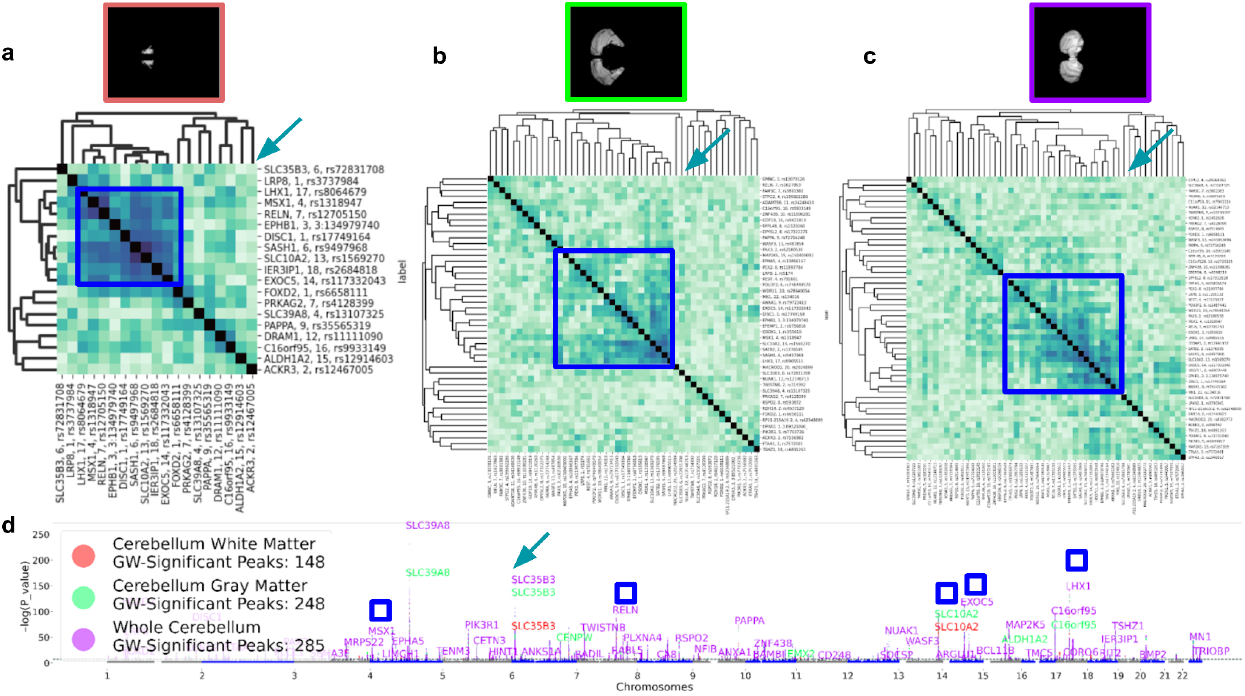
Cerebellar clustering. Hierarchical clustering in the latent space representation of lead SNPs reveals consistent structure in autoencoders trained on (a) only the white matter of the cerebellum, (b) only the gray matter of the cerebellum, and (c) the whole cerebellum. The blue square shows a clade containing SNPs from DISC1, RELN, SASH1 are similarly grouped in the 3 clusterings, while the turquoise arrow highlights how the SNP in SLC35B3 is consistently in a distant branch. (d) Manhattan plots of the 3 cerebellar latent spaces show distinct but overlapping structure.

In this way, working in the cross-modal ECG space we find two clusters corresponding to SNPs affecting the QT interval (SNPs associated with NOS1AP and KCNQ1) and SNPs related to the P-wave (SNPs associated with SCN10A and ALPK3), highlighted with green boxes in Figure 7 panel a. We also identify a large cluster corresponding to SNPs affecting multiple cardiac traits such as those associated with BAG3, SLC35F1, or KCND3 this larger group is mostly shared between the MRI and ECG spaces, as shown by the large blue square (with the notable exception of SCN10A).

**Figure 7:**
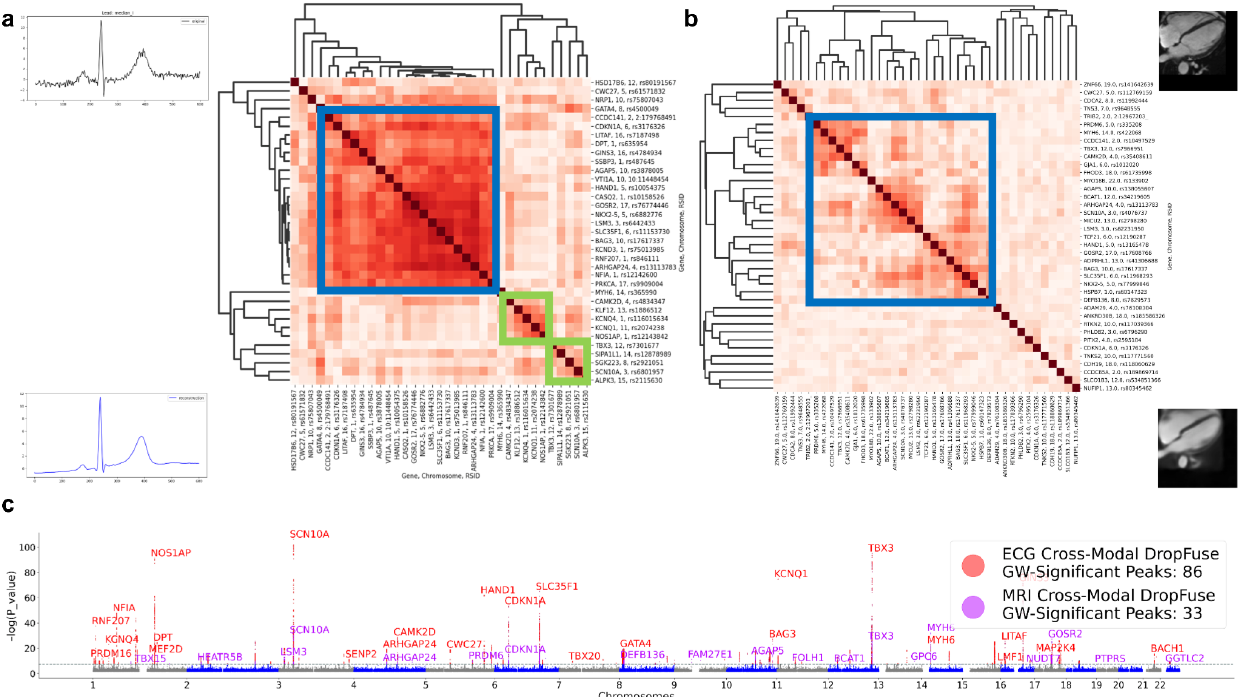
Cross-modal clustering. Hierarchical clustering in the cross-modal latent space of the (a) ECG and (b) cMRI. The blue square highlights a similar clade in both models with SNPs from CASQ2, BAG3, GOSR2, NKX2-5, while the green squares show two clades in the ECG latent space that are not seen in the MRI latent space. (c) Manhattan plots of the two cross-modal latent spaces.

## Limitations

There are situations where registration can introduce bias and distort associations. Anatomical atlases constructed in one population may not be appropriate for other populations with different demographics, ancestry or disease states. An advantage of the registration learning methods (VoxelMorph, DeepCycle, and DropFuse) over template-matching methods is that they don’t require prototypical individual(s) to be selected as reference. Still, learning methods are limited by the variation present in their training data. The UK Biobank, which we have studied here, is not representative of the world at large. Larger, more representative biobanks need to be built and existing models need to be inspected and/or corrected for potentially harmful bias. In our genetic analysis we removed known confounds with iterative nullspace projection after training our models, but this can also attenuate biological associations. Future methods might learn to automatically factorize signal and bias.

## Conclusion

In this work, we showed the importance of registration and fusion in learning holistic representations of physiological state. Many different registration methods across many imaging modalities consistently amplified biological signals. The best registration method for a given analysis will depend on many factors including computational resources, the availability, quality and applicability of anatomical atlases, as well as the level of interpretability desired. Moreover, some of the registration techniques discussed are complementary. For example, two modalities registered in space by VoxelMorph can later be cross-modally registered with DropFuse. Building up unified biologically informative latent spaces by combining many modalities and kinds of registration is an exciting avenue of future research.

