## Supplementary Material for "Genetic Architectures of Medical Images Revealed by Registration and Fusion of Multiple Modalities"

*Supplementary Video 1: deep_cycle_reconstructions.mov*
This video shows reconstructions from regularly spaced latent space samples from DeepCycle at left and from a single-parameter convolutional autoencoder without the inductive bias of periodicity on the right. DeepCycle has learned to encode the dynamics of the cardiac cycle, while the single parameter autoencoder without the circular inductive bias has encoded exposure value in the latent space.

*Supplementary Video 2: voxel_morph_cardiac_mri.mov*
This video shows the original cardiac MRI modality at left and at right the result of registration with voxel morph. Then the video compares the cardiacMRIs registered with a smoothing weight λ of 0.5 (left) 0.05 (right). The higher smoothing weight reduces high-frequency artifacts and was used for the results in the main paper.

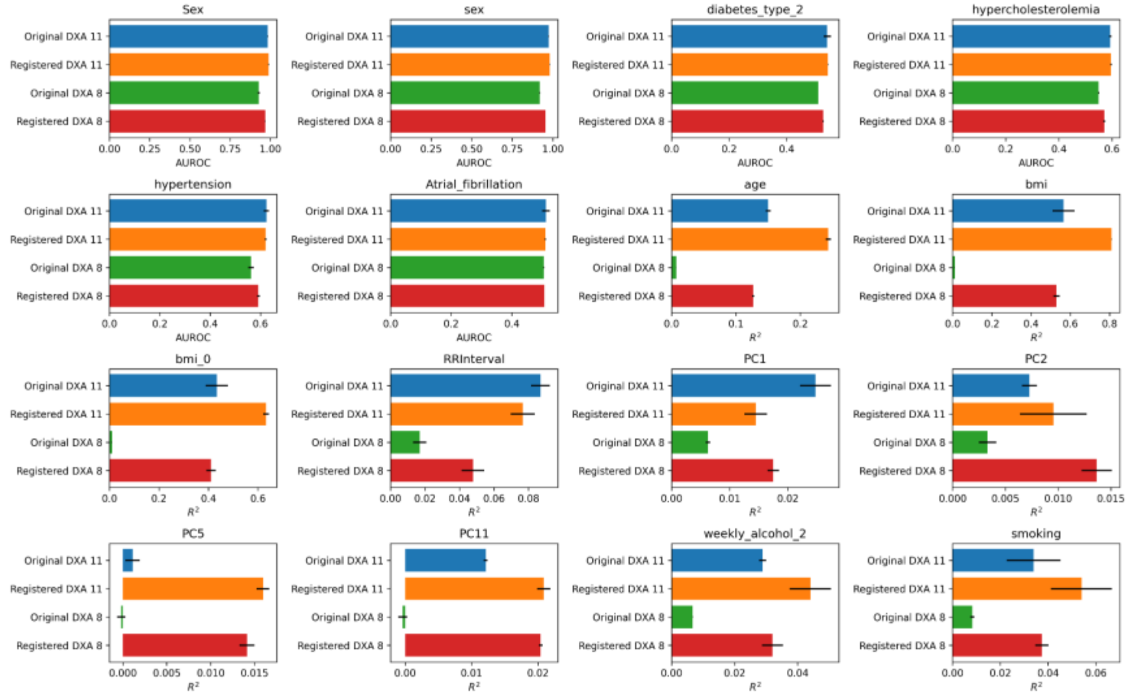


*Supplementary Figure 1: Continuous and categorical phenotypes predicted from latent spaces built from DXA series 8 and 11, in the original coordinate system and after DropFuse registration. Representations learned after registration encode more biological information. The two sex phenotypes are self-reported sex and genetically determined sex.*

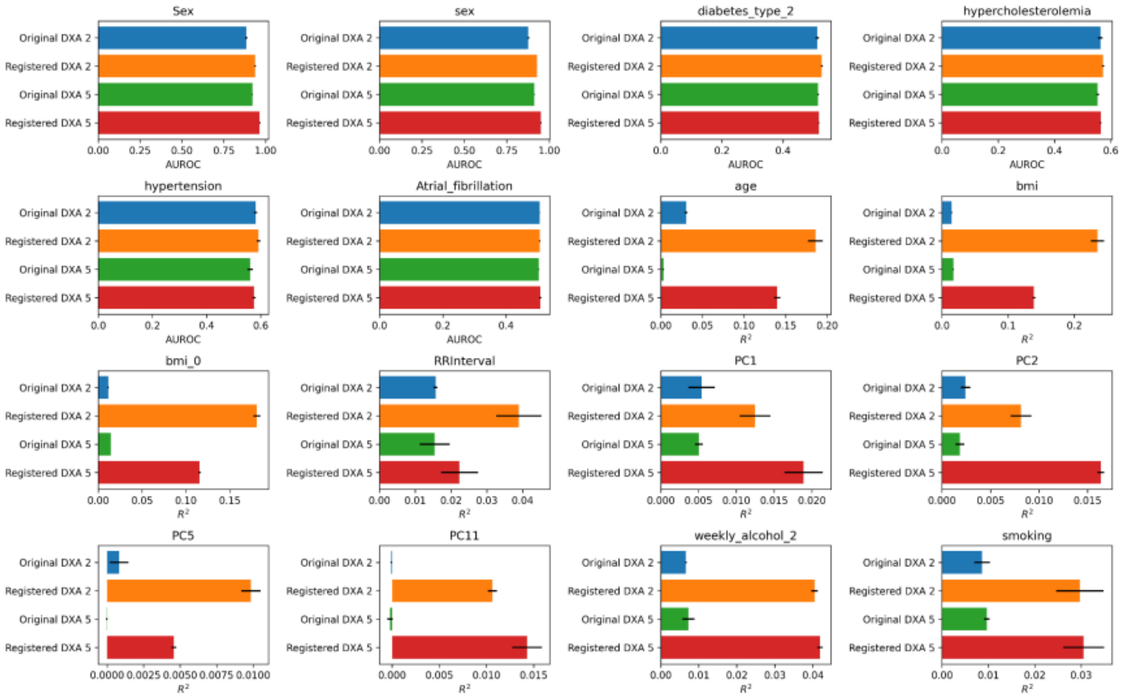


*Supplementary Figure 2: Continuous and categorical phenotypes predicted from latent spaces built from DXA series 2 and 5, in the original coordinate system and after registration. Representations learned after registration encode more biological information.*

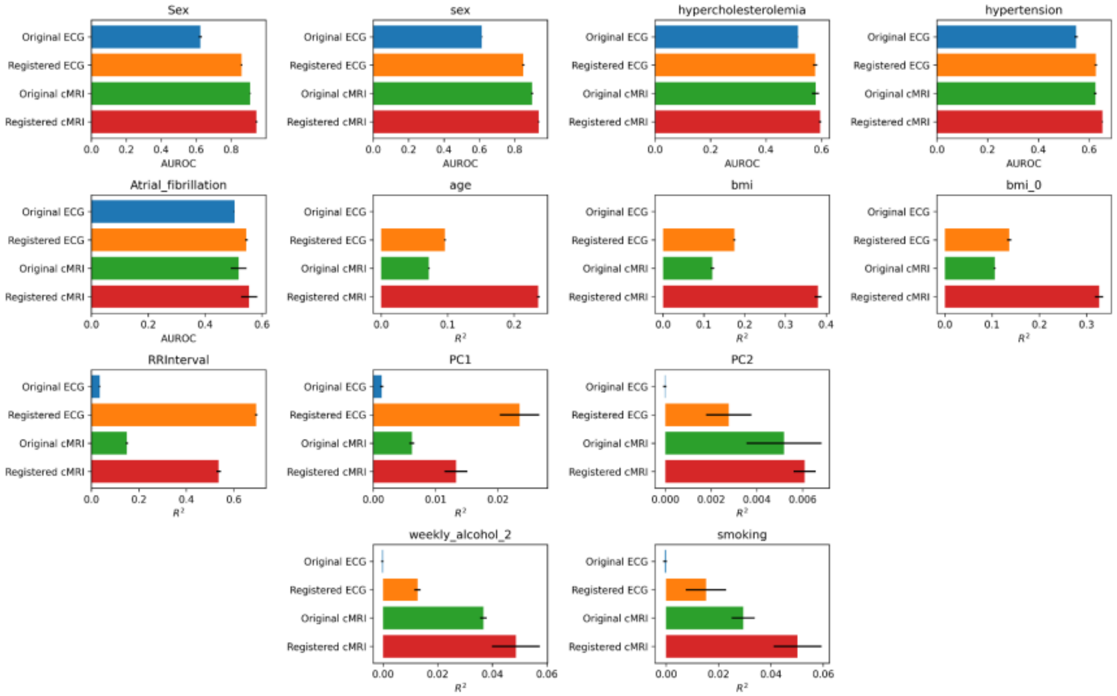


*Supplementary Figure 3: Continuous and categorical phenotypes predicted from latent spaces built from ECG and Cardiac MRI, in the original coordinate system and after DropFuse registration. Representations learned after registration encode more biological information.*

#### ECG PheWAS

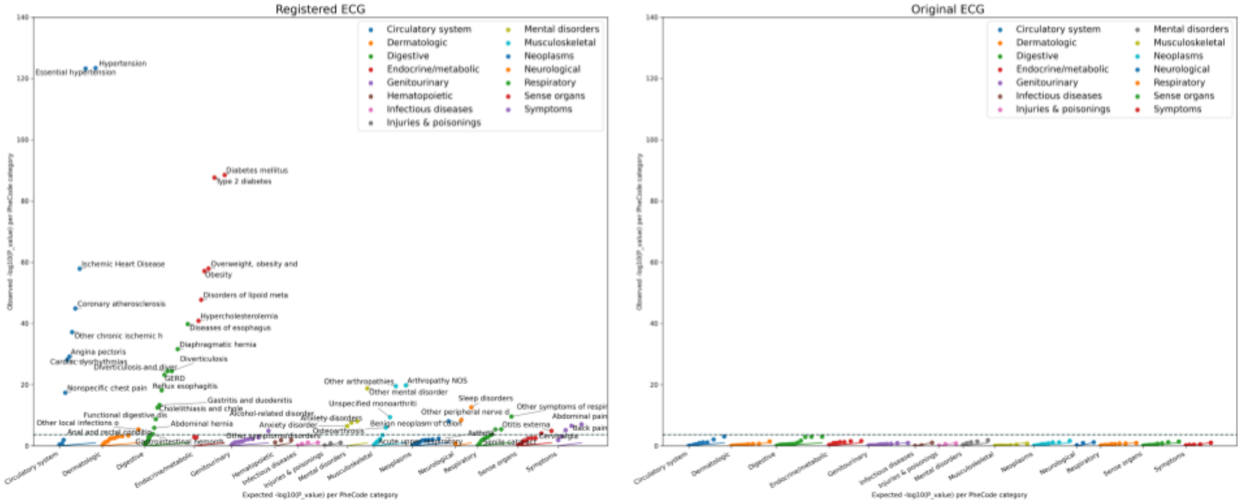


*Supplementary Figure 4: Phenome Wide Association Study of ECG latent spaces before and after registration. Registered modalities reveal more and stronger associations.*

#### Cardiac MRI PheWAS

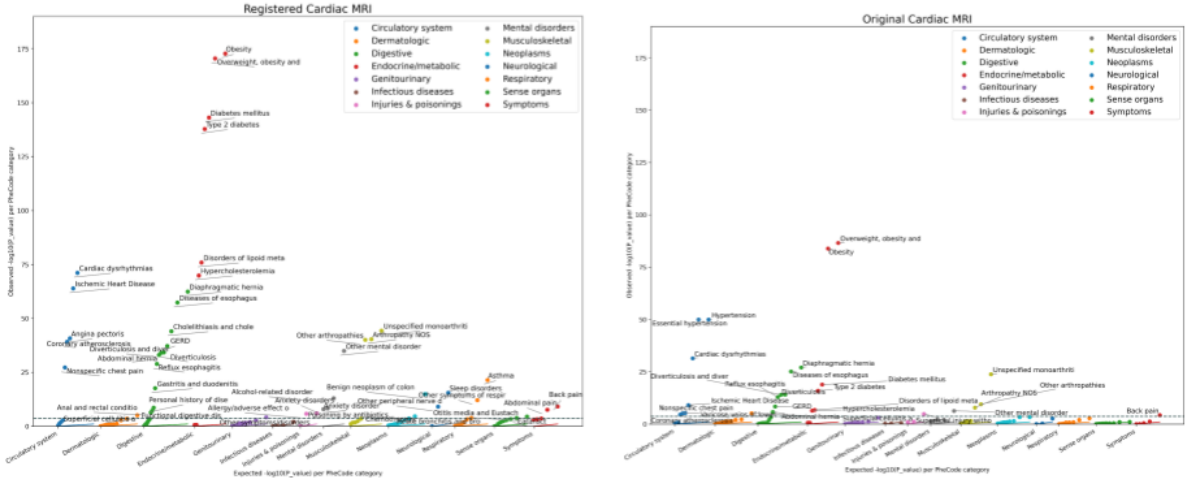


*Supplementary Figure 5: Phenome Wide Association Study of Cardiac MRI latent spaces before and after registration. Registered modalities reveal more and stronger associations.*

### Brain MRI PheWAS

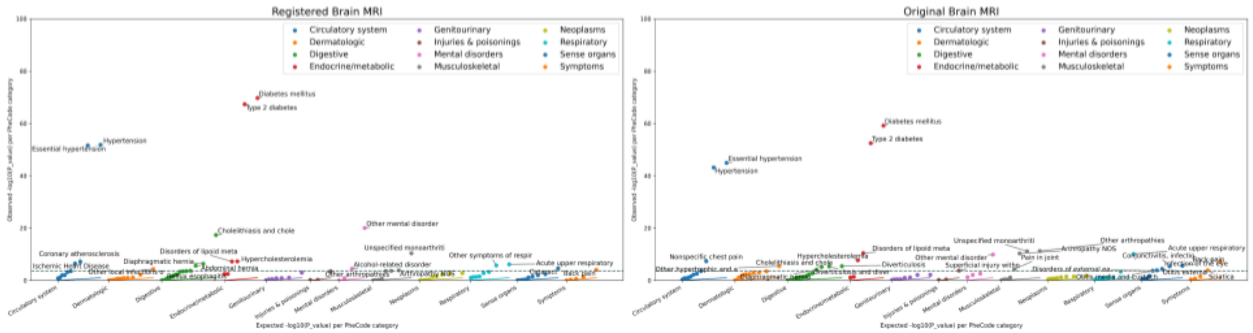


*Supplementary Figure 6: Phenome Wide Association Study of Brain MRI latent spaces before and after registration. Registered modalities reveal more and stronger associations.*

#### DeepCycle 1-parameter circular autoencoder learns the cardiac cycle from cMRI

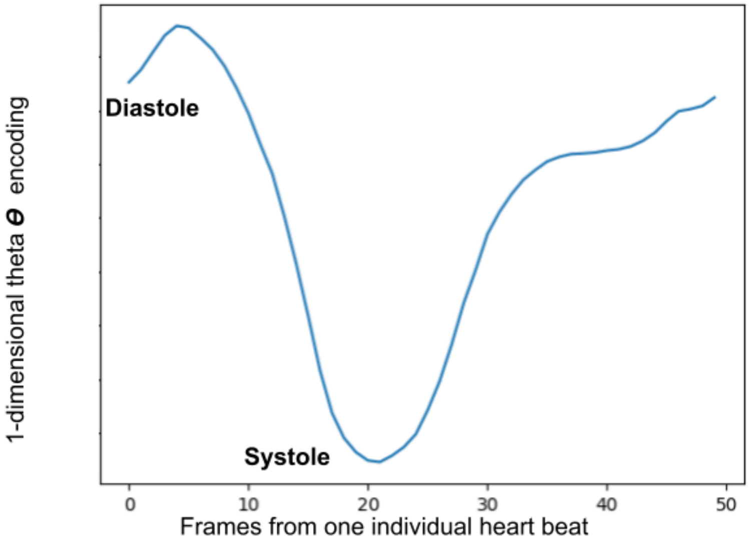


*Supplementary Figure 7: DeepCycle Cardiac MRI encoding from a single individual cardiac MRI of 50 frames spanning a single heartbeat.  The theta parameter encodes the dynamics of the cardiac cycle.*

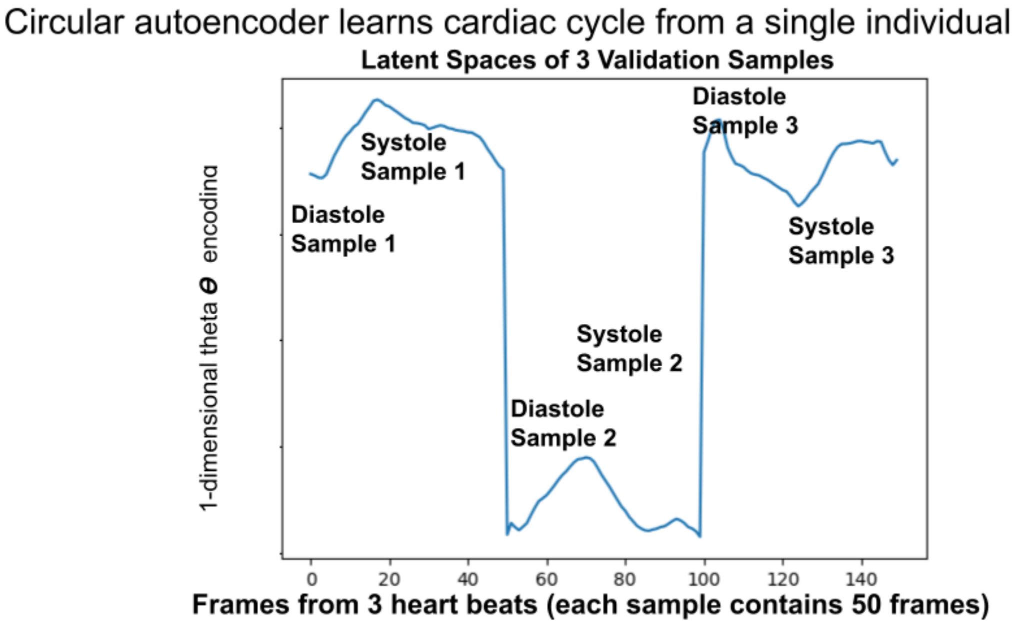


*Supplementary Figure 8: DeepCycle Cardiac MRI encodings from three individual's cardiac MRIs from the validation set. Each MRI contains 50 frames spanning a single heartbeat. Even when trained with just a single individual the theta parameter encodes the dynamics of the cardiac cycle on held-out individuals.*
